# Immune factor of bacterial origin protects ticks against host skin microbes

**DOI:** 10.1101/2020.04.10.036376

**Authors:** Beth M. Hayes, Atanas D. Radkov, Fauna Yarza, Sebastian Flores, Jungyun Kim, Ziyi Zhao, Katrina W. Lexa, Liron Marnin, Jacob Biboy, Victoria Bowcut, Waldemar Vollmer, Joao H. F. Pedra, Seemay Chou

**Author notes:** These authors contributed equally to this work (Beth M. Hayes, Atanas D. Radkov).

## Abstract

Hard ticks are blood-feeding arthropods that carry and transmit microbes to their vertebrate hosts^1^. Tick-borne disease cases have been on the rise over the last several decades, drawing much-needed attention to the molecular interplay between transmitted pathogens and their human hosts. However, far less is known about how ticks control their own microbes, which is critical for understanding how zoonotic transmission cycles persist. We previously found that ticks horizontally acquired an antimicrobial toxin gene from bacteria known as *<u>d</u>omesticated <u>a</u>midase <u>e</u>ffector 2* (*dae2)*^2^. Here we show that this effector from the tick vector *Ixodes scapularis* (Dae2^*Is*^) has structurally and biochemically diverged from ancestral bacterial representatives, expanding its antibacterial targeting range to include host skin microbes. Disruption of *dae2*^*Is*^ increases the burden of skin-associated staphylococci within *I. scapularis* and adversely affects tick fitness, suggesting resistance of host microbes may be important for the parasitic blood-feeding lifestyle. In contrast, Dae2^*Is*^ has no intrinsic ability to kill *Borrelia burgdorferi*, the tick-borne bacterium of Lyme disease. Our observations suggest that ticks have evolved to tolerate their own symbionts while resisting host skin commensals, which we discover are natural opportunistic pathogens of ticks. This work moves our understanding of vector biology beyond a human-centric view: just as tick commensals are pathogenic to humans, so too do our commensals pose a threat to ticks. These observations illuminate how a complex and mirrored set of interkingdom interactions between blood-feeding vectors, their hosts, and associated microbes can ultimately lead to disease.

## Main text

Blood-feeding (hematophagy) arose independently in at least six different arthropod lineages approximately 50-150 million years ago (MYA), coincident with the emergence of many immune and host-modulating genes predicted to support this hazardous lifestyle^3-5^. Hard ticks, such as the blacklegged tick *Ixodes scapularis*, have especially prolonged, continuous bloodmeals that can last over a week^1^, during which ticks must embed within the skin of their hosts and maintain an intimate, perilous attachment. *I. scapularis* ticks also occupy diverse environmental niches away from their host, where they spend the majority of their two-year lifecycles. We previously found that *I. scapularis* and other ticks acquired a potent antibacterial enzyme approximately 40 MYA by horizontal gene transfer of an interbacterial competition toxin gene from bacteria^2^. The timing and extraordinary interkingdom journey of this <u>d</u>omesticated <u>a</u>midase <u>e</u>ffector 2 (*dae2*) suggests that antibacterial defense is important for the remarkable hematophagous parasitism of hard ticks. However, the microbes driving *dae2* acquisition and tick immune expansion are not well defined.

*I. scapularis* innate immunity has likely evolved to resist bacteria that pose a threat to tick fitness. The microbiota of *I. scapularis* ticks are dominated by a few stably-associated symbionts, such as *Rickettsia spp*. and the causative agent of Lyme disease *Borrelia burgdorferi*^6-11^, which can be transmitted to humans through the tick bite. Although these microbes are disease-causing human pathogens, the nature of their relationships with vector tick hosts is less clear. Acquisition of *B. burgdorferi* during feeding activates expression of numerous tick immune genes, including *dae2* from *I. scapularis* (*dae2*^*Is*^)^12^. We previously found that *dae2*^*Is*^ modestly limits *B. burgdorferi* levels in *I. scapularis*^2^. Paradoxically, *B. burgdorferi* is a stably-associated tick microbe that is not cleared by *I. scapularis* immunity and does not appear to negatively impact tick fitness^13,14^. These observations suggest that Dae2 may promote tick tolerance of *B. burgdorferi* by preventing excessive proliferation in the vector, similar to what has been observed for host–commensal interactions in other systems^15^. Alternatively, or in addition, Dae2 enzymes may function to resist and protect against less stably-associated microbes that are true pathogens of ticks. Indeed, the highly host-dependent status of vector-borne microbes as both symbionts and pathogens is at the very crux of how the zoonotic diseases persist. Defining the role of immune effectors, such as Dae2, would provide a window into how such complex, multi-kingdom interactions between ticks, their hosts, and microbes are maintained^16,17^.

We posit that the ancestral function of *dae2* genes provides some clues as to its biological role in ticks. The *dae2* genes originated from horizontally co-opted bacterial <u>t</u>ype VI secretion (T6S) <u>a</u>midase <u>e</u>ffector 2 (*tae2*) genes, which encode lytic cell wall-degrading enzymes. The *tae* genes encode a family of interbacterial competition toxins that bacteria use to break down bacterial cell wall peptidoglycan (PG) specifically from rival Gram-negative bacteria. The Tae enzymes are delivered by specialized T6SS injection machinery into the membrane-bound cell wall compartment (periplasm) of recipient Gram-negative bacteria^18,19^, resulting in osmotic stress and cell lysis. In contrast, the eukaryotic *dae* genes are not associated with any specialized secretion apparatus. Although Dae enzymes have retained the ability to degrade PG, the eukaryotic representatives are not intrinsically capable of traversing the protective outer membrane of Gram-negative bacteria^2^. Thus, we hypothesize that Dae effectors may act against other microbial groups or function via alternative mechanisms. Furthermore, given that evolutionary and functional divergence of domesticated genes is common in cases of horizontal gene transfer between highly disparate organisms, we hypothesize that Dae2 in ticks may act through mechanisms distinct from bacterial Tae toxins.

To investigate, we conducted a sequence-based comparison of the domesticated *dae2* genes of ticks and mites with ancestral bacterial *tae2* homologs. This showed several regions of variance between these groupings (Extended Data Fig. 1a and 1b). Although we were unable to find definitive evidence of positive or negative selection^2^, these variances were unique to eukaryotic representatives and highly suggestive of functional differences. After crystallization trails with several Tae2/Dae2 family members, we were able to determine the high-resolution X-ray crystal structure of one Tae2/Dae2 family member, Tae2 from *Salmonella enterica* Typhi (Tae2^*St*^) at a resolution of 2.05 Å (Fig. 1a, Extended Data Table 1, PDB ID: 6WIN). Tae2^*St*^ adopts an amidase fold conserved across papain-like proteases and most similar to endolysin LysK from staphylococcal phage K (Extended Data Fig. 2)^20^. From this structure, we generated threaded homology models of remaining family members at a confidence interval (>90% confidence) that allowed high probability fold prediction^21^. Alignment of the Tae2^*St*^ structure and a Dae2^*Is*^ homology model (112 aligned residues; 32% identity) pointed to several substantial structural divergences between eukaryotic and prokaryotic subgroups that correlated with our sequence-based analyses, mapping the three major regions of sequence divergence mapped to the loops (L1-L3) that abut the catalytic channel (Fig. 1b). This comparison also revealed substantial differences in the topology and electrostatic properties of a deep hydrophobic substrate-binding groove that extends from the active site (Fig. 1c and 1d). The catalytic residues themselves are more accessible in Dae2^*Is*^ than in Tae2^*St*^, with a predicted pore behind the catalytic cysteine of Dae2^*Is*^. Surface conservation profiles (Extended Data Fig. 3) further suggested that these divergent properties are conserved within Tae2 and Dae2 subgroups, respectively. Importantly, the modeled differences in loop structures, the substrate-binding groove and the catalytic channel led us to hypothesize that the substrate specificity of Dae2 could either be distinct from or expanded compared to the bacterial enzymes.

**Figure 1.**
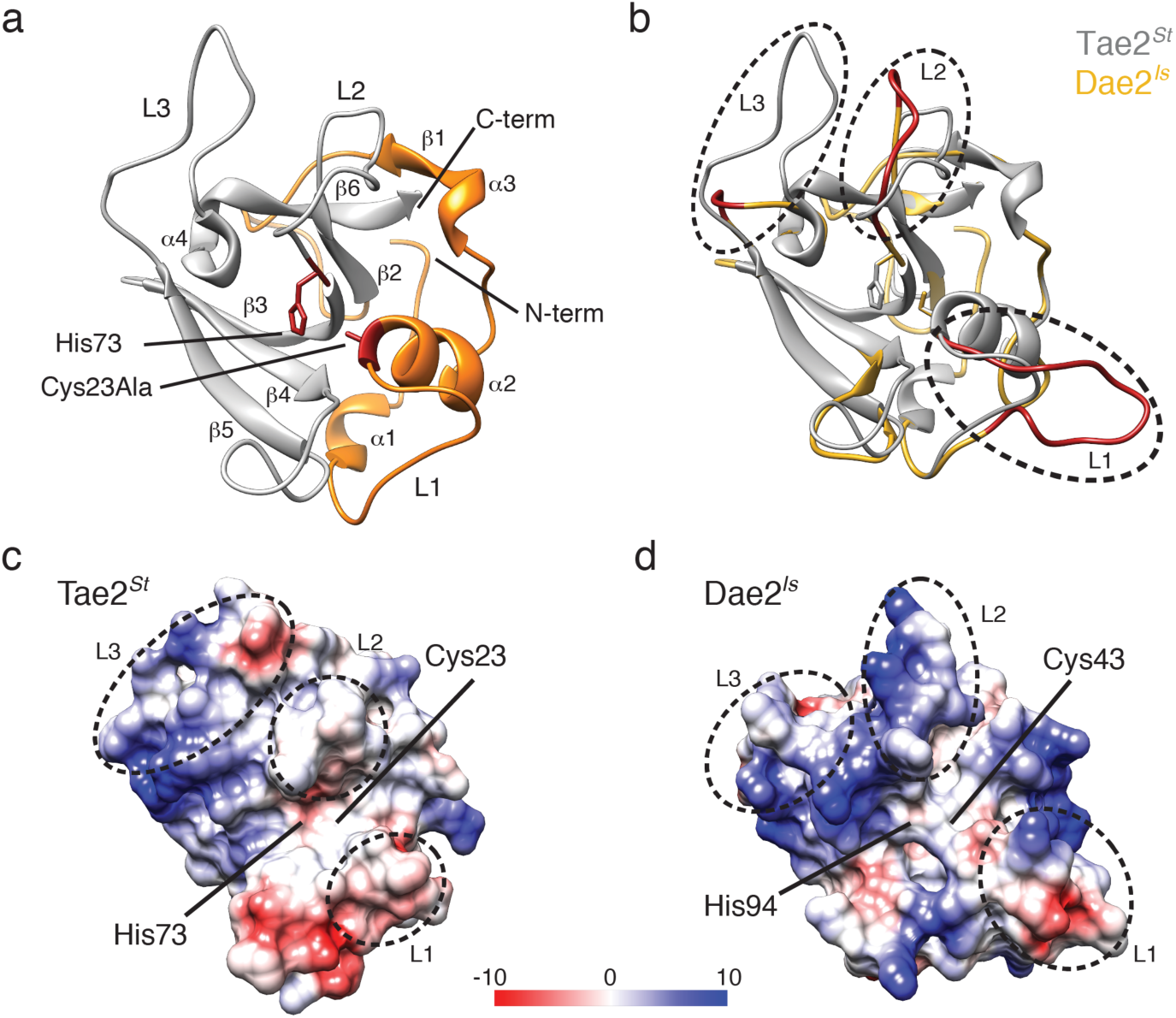
Tae2 and Dae2 enzymes have divergent structural features. **a**, Ribbon diagram Tae2 from *Salmonella enterica* Typhi (Tae2^*St*^). Canonical papain-like secondary structures are labeled (α1-4, β1-6, Loop1-3 (L1-3)), and subdomains are colored (gray, orange). Catalytic residues are denoted (labeled His 73 and Cys23Ala, red). **b**, Structural alignment of Tae2^*St*^ (gray) and a Tae2-based model of Dae2^*Is*^ (golden). Loops 1, 2, and 3 are colored red in Dae2^*Is*^. **c, d**, Electrostatic charges in surfaces proximal to the catalytic residues for Tae2^*St*^ and Dae2^*Is*^ (scale is in kcal mol^-1^ *e*^-1^). The loops are also boxed in the sequence alignment presented in Extended Data Fig. 1.

**Figure 2.**
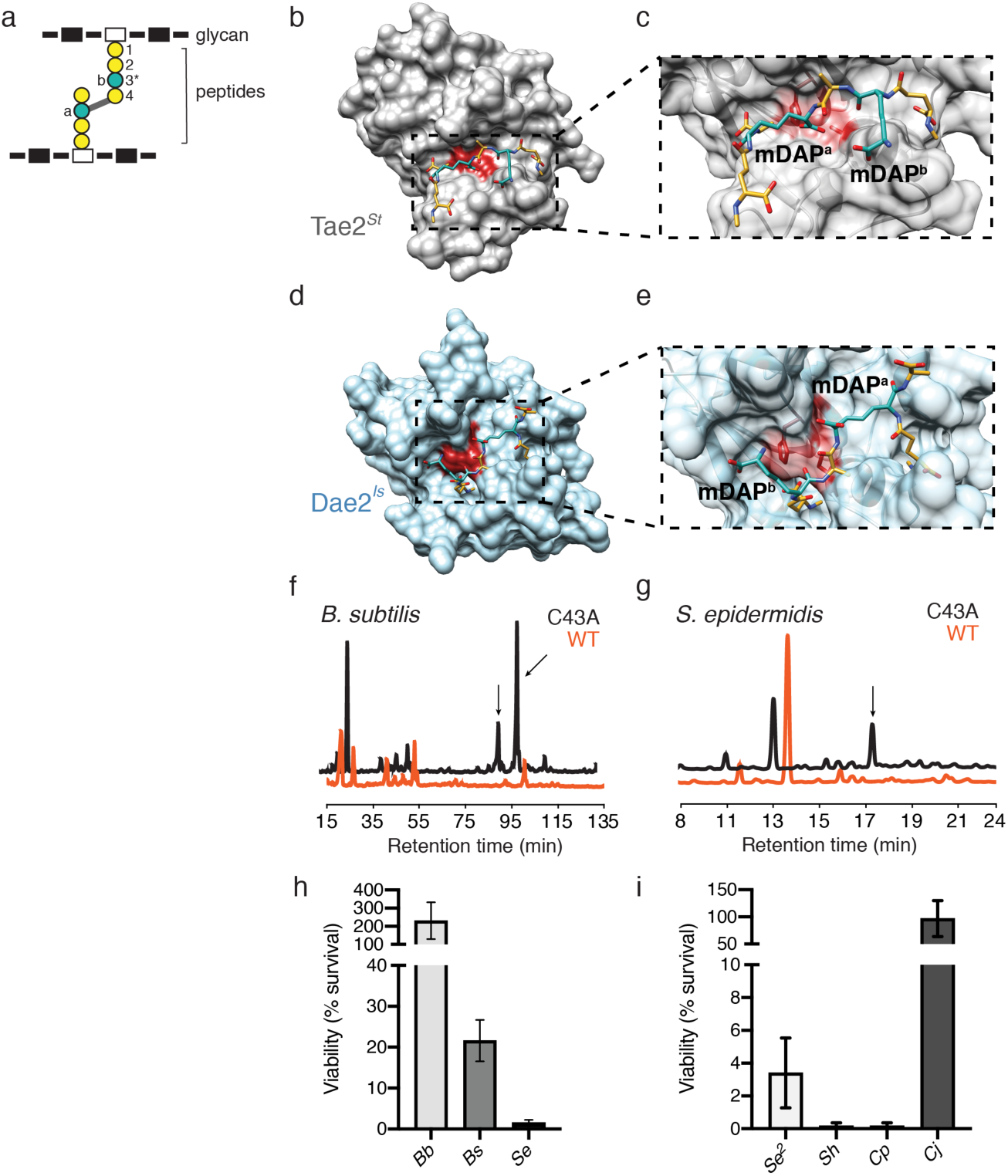
Enzymatic and antibacterial specificity of Tae2 and Dae2. **a**, Cartoon schematic of bacterial peptidoglycan (PG) structure with glycans (rectangles), peptides (circles), and cross-link position (gray line) shown. Residue positions in PG peptide stem are labeled (1-4) with key variable third position (3*) colored in teal. **b**, Surface representation of Tae2^*St*^ ligand docking using a stem peptide substrate. Both *m*DAP residues in the stem peptide are colored in teal. **c**, Close-up of the Tae2^*St*^ active site and docked substrate. The scissile amide bond is between *m*DAP^a^ and the adjacent D-ala (position 4), proximal to the active site Cys23 residue. **d**, Surface representation of Dae2^*Is*^ ligand docking, using the same substrate as for Tae2^*St*^ in C. **e**, Close-up of the Dae2^*Is*^ active site and docked substrate. The scissile bond is similarly situated between *m*DAP^a^ and the adjacent D-ala, proximal to the active site Cys43 residue. Notice the substrate directionality being *m*DAP^b^ – D-ala – *m*DAP^a^, compared to the inverse for Tae2^*St*^, *m*DAP^a^ – D-ala – *m*DAP^b^. HPLC chromatograms of PG from *B. subtilis* **f**, and *S. epidermidis*. **g**, after enzymatic degradation with Dae2^*Is*^ wt (orange) and its catalytically inactive mutant (C43A, black). Arrows indicate the substrate peaks. Dae2^*Is*^ lysis assays of **h**, Gram-negative *Borrelia burgdorferi* strain S9 (*Bb*), Gram-positive *Bacillus subtilis* 168 (*Bs*), and Gram-positive *Staphylococcus epidermidis* BCM060 (Se) and **i**, Gram-positive skin associated bacteria *S. epidermidis* strain SK135 (*Se*^2^), *S. hominis* strain SK119 (*Sh*), *Corynebacterium propinquum* DSM44285 (*Cp*) and *C. jeikeium* DSM7171 (*Cj*). Results are reported as percent survival with Dae2^*Is*^ wt normalized to its catalytic mutant C43A.

**Figure 3.**
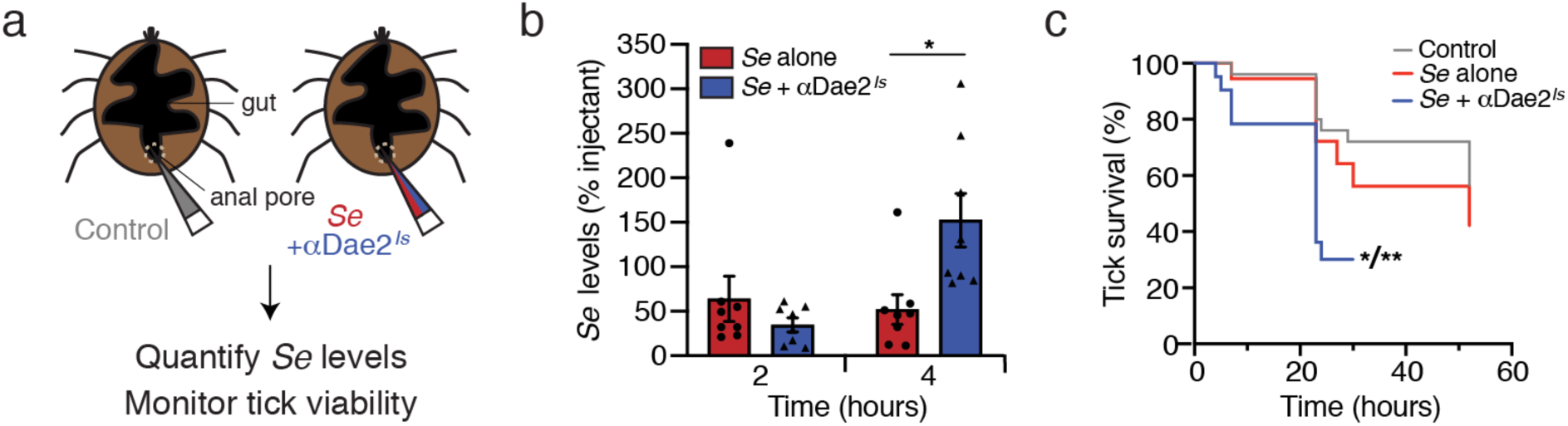
Dae2-dependent effects on *I. scapularis–S. epidermidis* interactions *in vivo*. **a**, Schematic of experimental setup. *I. scapularis* nymphs were microinjected into the anal pore with a control antibody (grey, left), live *S. epidermidis* alone (red, right), or live *S. epidermidis* and αDae2^*Is*^ together (blue, right). Ticks were then monitored for levels of *S. epidermidis* and tick survival. **b**, CFU levels of *S. epidermidis* alone (red) or together with αDae2^*Is*^ antibody to neutralize the lytic activity of Dae2^*Is*^ (blue) at 2 and 4 hours post injection. Results are shown as percentage of CFU at time 0 (injectant). *p<0.05 based on paired t-test. **c**, Tick survival after microinjection of αDae2^*Is*^ control (grey), *S. epidermidis* alone (red), or *S. epidermidis* together with αDae2^*Is*^ antibody to neutralize the lytic activity of Dae2^*Is*^ (blue). *p<0.05 compared to *S. epidermidis* alone and **p<0.01 compared to antibody alone based on Log-rank (Mantel-Cox) test.

Divergent Dae2 specificity could have important biological implications for tick immunity. Given that cell wall composition varies considerably between bacterial taxa^22^, enzyme selectivity for PG types directly dictates which microbes can be inhibited. Of note, Gram-negative bacteria incorporate *meso-*diaminopimelic acid (*m*DAP) at the third position in the peptide stem (Fig. 2a)^22^, whereas bacteria from the class Bacillus have an amidated *m*DAP and other Gram-positive species have a lysine^22^. We previously found both Tae2 and Dae2 can hydrolyze Gram-negative cell walls^2^ (Extended Data Fig. 4), and a Tae toxin from *Pseudomonas aeruginosa* is highly selective for *m*DAP-containing Gram-negative PG^23^. We employed an *in silico* approach to further probe specificity by computationally docking a fragment of crosslinked *m*DAP PG onto Tae2^*St*^ and Dae2^*Is*^ (Fig. 2b-e). Docked PG adopted highly dissimilar poses between the two structures. We calculated an overall narrower cleft on Tae2^*St*^ with 852.9 Å^2^ active site surface area versus 1086.0 Å^2^ for Dae2^*Is*^, which also has an additional deep binding pocket. Upon binding, the cross-linked PG interacted with Dae2^*Is*^ across a larger area of the catalytic site (624.9 Å^2^) compared to Tae2^*St*^ (450.2 Å^2^). Our modeling and metadynamics simulations predicted a flipped PG orientation mediated through different residue interactions at the enzyme–substrate binding interface (Extended Data Fig. 3c and 3d) as well as tighter Dae2– PG binding. We hypothesize that altered binding and positioning of the PG substrate could lead to an enzyme specificity profile for Dae2 that is distinct from their bacterial antecedents. The widened catalytic channel of Dae2 leads us to predict that the eukaryotic enzymes can act on a broader range PG structural groups relative to Tae2 toxins, widening the range of targeted bacteria.

**Figure 4.**
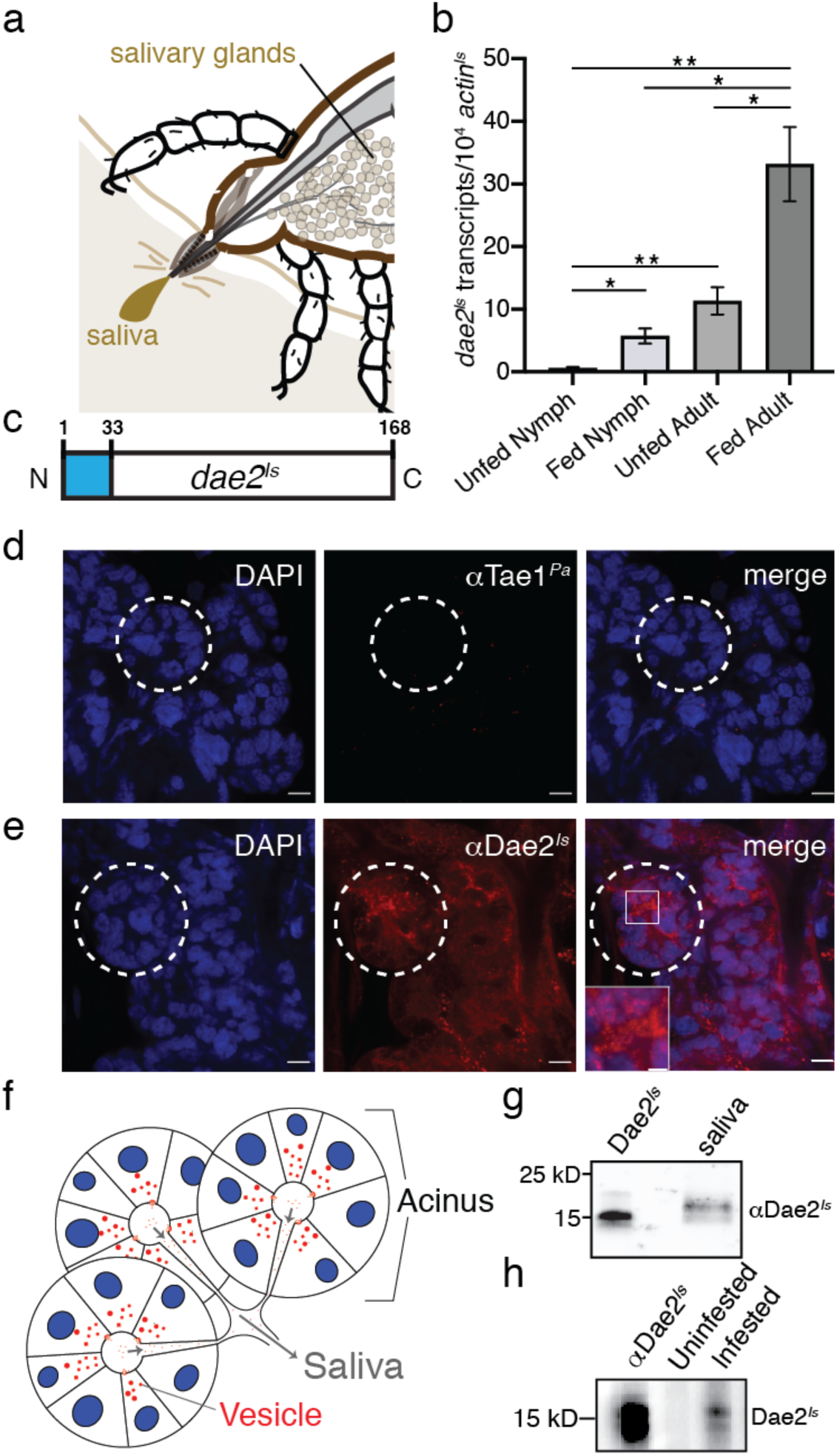
Dae2^*Is*^ is upregulated and found in saliva of ticks during feeding. **a**, Schematic of the tick at the bite-site interface showing the salivary glands. The salivary glands produce the saliva that is injected into the host. **b**, *dae2*^*Is*^ expression (normalized against *actin*^*Is*^ transcripts) before (unfed) and after feeding (fed) on mice for nymph and adult ticks. * *p*< 0.05, ** *p*<0.01. **c**, Diagram of the *dae2*^*Is*^ gene structure showing the predicted N-terminal signal sequence from nucleotide position 1-33 (blue). **d, e**, Confocal images of *I. scapularis* salivary glands from adult females. Whole mounts were immunostained with DAPI (left panels, blue), control rabbit antibody against Tae1^*Pa*^ (d, middle, red) or rabbit αDae2^*Is*^ (e, middle, red). Scale bars= 15µm. Selected acini are marked (dashed white lines). A zoomed-in region is shown (box). **f**, Cartoon representation of salivary gland acini. Nuclei (blue), vesicles (red), and saliva flow routes (grey arrows) are shown. **g**, Western blot of collected saliva from partially fed *I. scapularis* adult females demonstrating presence of Dae2^*Is*^ in saliva. First lane has recombinant Dae2^*Is*^ as a positive control for rabbit αDae2^*Is*^ reactivity. **h**, Western blot showing reactivity of mouse serum to Dae2^*Is*^. Recombinant Dae2^*Is*^ protein was probed with mouse serum before (uninfested) or after *I. scapularis* feeding (infested). Rabbit αDae2^*Is*^ was used as positive control for reactivity (first lane).

We experimentally tested our specificity model by comprehensively assessing the *in vitro* substrate specificity of Dae2^*Is*^ and Tae2^*St*^ enzymes. Purified recombinant proteins were tested against a set of PG sacculi prepared from bacteria that we and others have found associated with field *I. scapularis* ticks: two environmental species (*Escherichia coli*, Gram-negative; *Bacillus subtilis*, Gram-positive) and a mammalian skin-associated bacterium (*Staphylococcus epidermidis*, Gram-positive)^8,24^. Analysis of PG degradation by high performance liquid chromatography (HPLC) showed that the bacterial Tae2^*St*^-dependent hydrolysis was restricted to Gram-negative cell walls (Extended Data Fig. 4a-c), consistent with the defined role Tae2 toxins play in T6S interbacterial competition between Gram-negative bacteria. On the other hand, Dae2^*Is*^ hydrolyzed all PG sacculi types tested (Fig. 2f-g), including those from lysine-containing Gram-positive cell walls, which has not been previously reported for Tae enzymes. Expansion of Dae2^*Is*^ activity to Gram-positive *S. epidermidis* PG was especially remarkable in light of how chemically divergent its cell wall composition is from Gram-negative PG. Our biochemical observations open up the exciting possibility that Dae2^*Is*^ has evolved to lyse a wider range of microbes, which may include skin-associated microbes found at the tick–host feeding interface.

To examine whether our Dae2^*Is*^ *in vitro* specificity profile reflects broadened antibacterial activity, we next assayed Dae2^*Is*^-dependent effects on bacterial cell viability. We carried out bacterial killing assays at a physiologically relevant concentration of 2 μM Dae2^*Is*^, which was determined through quantification of levels in dissected *I. scapularis* salivary glands (Extended Data Fig. 5). *E. coli* is intrinsically resistant to Dae2^*Is*^, which is likely due to the inability of Dae2^*Is*^ to penetrate its outer membrane and access the PG^2^. We similarly found that Dae2^*Is*^ was not able to kill the tick-borne spirochete, *B. burgdorferi* (Fig. 2h), likely because it too has an outer membrane, although it is highly distinct from that of *E. coli*^25,26^. However, given that Dae2^*Is*^ can degrade purified *B. burgdorferi* cell wall fragments^2^, we cannot rule out the possibility that Dae2^*Is*^ acts in synergy with an additional host factor or process *B. burgdorferi* PG to modulate canonical innate immune signaling pathways encoded by *I. scapularis*^27,28^. In stark contrast, Dae2^*Is*^ was highly efficient at directly killing *S. epidermidis*, a human skin commensal, and moderately efficient at killing *B. subtilis*, a common environmental bacterium (Fig. 2h). These dramatic differences lead us to hypothesize that human skin microbes, which are frequently encountered by feeding ticks, are targeted by Dae2^*Is*^. We further tested Dae2^*Is*^ against three additional skin commensal species: another *S. epidermidis* strain, *S. hominis* and *Corynebacterium propiquum*, and observed immediate lysis of all (Fig. 2i). Although related to *C. propiquum* and associated with human skin and mucus membranes, we did not observe killing of *C. jeikeium* by Dae2^*Is*^ (Fig. 2i), suggesting that Dae2^*Is*^ specificity is broad but not indiscriminate. It is possible that this more invasive strain of *Corynebacterium* has evolved mechanisms of resistance to cell wall targeting enzymes^29^. Together, our analyses led to two major conclusions: expanded Dae2 specificity allows the eukaryotic enzymes to target a broader set of microbes than the prokaryotic Tae2 enzymes, and Dae2 kills common mammalian skin commensals with incredible potency.

**Figure 5.**
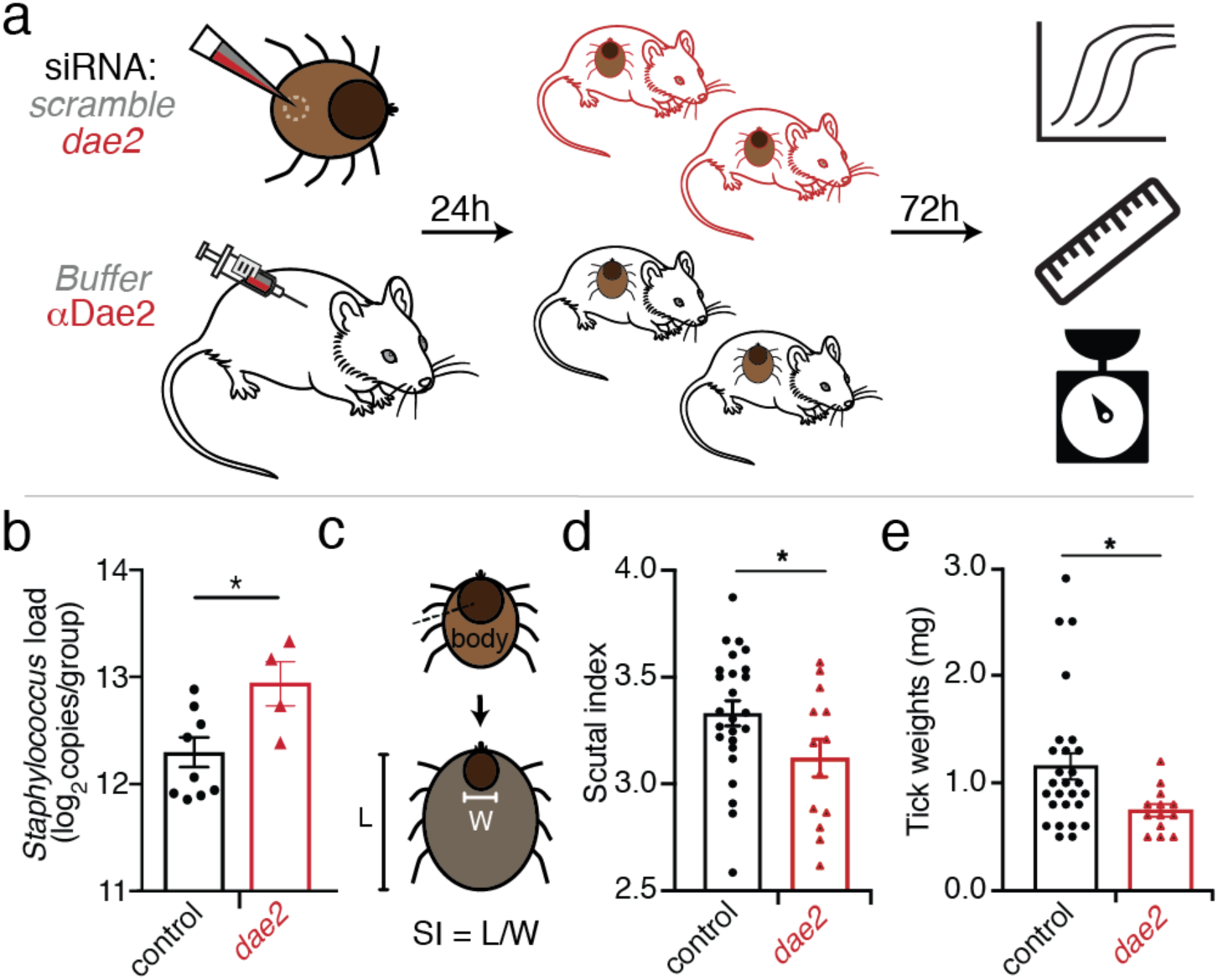
Dae2 activity affects tick fitness during feeding on mice. **a**, Schematic of *in vivo dae2*^*Is*^ knockdown experiment. Mice were injected with sterile saline control (grey) or 2.5mg Rabbit αDae2^*Is*^ antibody and *I. scapularis* nymphs were microinjected with siRNAs targeting *dae2*^*Is*^ (red) or scrambled controls (grey). After 24 hour recovery, *dae2*^*Is*^ injected and scrambled injected ticks were placed on αDae2^*Is*^ antibody injected mice or control mice, respectively. Ticks were forcibly detached at 3 days and assessed by qPCR, scutal index, and weight. **b**, *Staphylococcus* load was detected in DNA isolated from pooled *dae2*^*Is*^ knockdown (red) or control ticks (black) by qPCR with primers designed to target all species from the genus *Staphylococcus*. Results are presented as the average log_2_ *Staphylococcus* per tick. **c**, Diagram of how scutal index (SI) changes during feeding. Scutal index is the ratio of the length of the entire tick body to the width of the scutum^33^. **d**, Scutal index and **e**, tick weight of individual ticks from experiment detailed in Fig. 5b. *p<0.05 based on student’s t-test. Tick length and scutal width were measured from pictures with ImageJ. Weight was measured with an analytical balance with an error of +/- 0.1mg.

Hard ticks stay attached to the host skin dermis for days up to over a week, and this sustained interaction likely leads to intimate interactions between feeding ticks and microbial commensals of host skin. Our biochemical observations potentiate the model that Dae2^*Is*^ has the enzymatic capacity to kill bacterial groups commonly found at this critical tick–host bloodmeal interface. To evaluate this model, we first assessed whether physiological levels of Dae2^*Is*^ can inhibit *S. epidermidis* levels *in vivo* through artificial challenge assays using live nymphal *I. scapularis* ticks (Fig. 3a). Midguts of individual nymphs were infected with 10^2^ log-phase *S. epidermidis* cells through needle-based microinjections into their anal pores. At 4 hours post-injection, we compared levels of *S. epidermidis* from ticks with and without antibody-based biochemical inhibition of Dae2^*Is*^ in their guts. For these experiments, we raised and affinity-purified a rabbit antibody against recombinant Dae2^*Is*^ (αDae2^*Is*^). αDae2^*Is*^ specificity was validated by Western blot analysis (Extended Data Fig. 6a), and inhibitory function was confirmed through *in vitro* Dae2^*Is*^ lysis assays (Extended Data Fig. 6b). To deplete Dae2^*Is*^ activity within the tick midgut for our challenge assays, ticks were co-injected with *S. epidermidis* and our neutralizing αDae2^*Is*^. Inhibition of Dae2^*Is*^ led to higher levels of *S. epidermidis*, suggesting Dae2^*Is*^ has the capacity to directly antagonize *S. epidermidis in vivo* (Fig. 3b) by killing or inhibiting cell growth. Strikingly, we also observed a notable concomitant increase in the death of the ticks themselves in the *S. epidermidis*+αDae2^*Is*^ group (Fig. 3c) compared to *S. epidermidis* alone and control groups. These effects represent a clear fitness cost for ticks harboring high levels of *S. epidermidis* and are highly suggestive of an antagonistic, pathogenic relationship. The negative interactions between ticks and *S. epidermidis* is in keeping with the observation that this commensal is a highly adapted microbial partner of mammalian skin, a host environment that is very distinct from the tick and its gut. These experiments point to a potential role for gut Dae2^*Is*^ in resisting host skin commensals that could enter the tick during feeding.

In addition to Dae2^*Is*^ activity in the gut, we reason that the enzyme may play a second, additional role at the host–tick interface to also limit microbial entry. This prediction stems from our previous observation that Dae2^*Is*^ is highly enriched in *I. scapularis* salivary glands^2^. Moreover, it is well documented that ticks and other hematophagous arthropods deliver a variety of secreted molecules, such as antimicrobial and host-modulating effectors, to host bite sites via saliva to ensure successful bloodmeal completion (Fig. 4a)^5,30^. Such a function for Dae2^*Is*^ would be contingent on its presence in tick saliva over the course of a bloodmeal. Thus, we tracked its expression and precise subcellular localization during feeding. We quantified *dae2*^*Is*^ transcripts throughout two feeding cycles and found dramatic upregulation of *dae2*^*Is*^ expression in both nymph and adult ticks following bloodmeal consumption (Fig. 4b). We imaged Dae2^*Is*^ distribution in *I. scapularis* salivary glands by confocal microscopy and found that it localizes to vesicle-like puncta (Fig. 4d-e; Extended Data Fig. 7). Vesicle-mediated delivery of effector molecules to hosts has been observed in other blood-feeding vector arthropods^31^. The *dae2*^*Is*^ N-terminus encodes a predicted signal peptide (Fig. 4c)^2^; thus, we posited that Dae2^*Is*^ can be secreted out of tick salivary cells into saliva, enabling access to microbes near the bite site. To test this, we used a glass capillary to harvest saliva produced by live, partially-fed *I. scapularis* ticks (Fig. 4f). Immunoblot analysis of the extracted sample showed that Dae2^*Is*^ is indeed secreted into the outgoing saliva. We observed a slight increase in protein size relative to recombinant protein, suggestive of posttranslational modifications in the tick. We also tested whether Dae2^*Is*^ from saliva may physically enter the bloodmeal host by examining immunoreactivity of blood sera collected from mice that had been fed on by *I. scapularis* ticks (infested). Sera from mice hosts showed immunoreactivity to Dae2^*Is*^ after a single round of tick infestation but not for mice that had never been exposed to ticks (Fig. 4g). These results indicate that the tick injects Dae2^*Is*^ via saliva into the bite site where it is recognized as an antigen by the murine immune system. The timing and localization of Dae2^*Is*^ in tick saliva strongly suggests that this effector has antibacterial functions at multiple sites during feeding, which would impose multiple defense barriers against infectious microbes that ticks are likely to encounter while feeding.

To ask whether Dae2^*Is*^ activity in ticks contributes to resistance against host commensals during feeding, we next tested whether genetic disruption of *dae2*^*Is*^ in *I. scapularis* affects levels of the host skin-associated bacteria in nymphal ticks. We knocked down *dae2*^*Is*^ expression through microinjection-based RNA interference (RNAi) prior to feeding *I. scapularis* nymphs on mice (Fig. 5a). We also blocked pre-existing Dae2^*Is*^ protein in tick midguts or saliva by immunizing mouse bloodmeal hosts with our inhibitory αDae2^*Is*^ 24 hours before tick feeding (Fig. 5a). We found that αDae2^*Is*^ remained stable and detectable in mouse sera up to at least two weeks post-immunization (Extended Data Fig. 8), allowing us to inhibit Dae2^*Is*^ activity throughout the entirety of a tick bloodmeal. We retrieved ticks during the fast-feeding phase of their bloodmeals (day 3) and measured levels of host-associated bacteria present in ticks by qPCR, using genus-specific primers that reflect the diversity of *Staphylococcus* commensal strains colonizing mouse skin (Extended Data Table 2). Ticks where Dae2^*Is*^ activity was significantly reduced or absent supported higher levels of *Staphylococcus* bacteria than the control ticks (Fig. 5b). Our *ex vivo* challenge experiments suggested that elevated levels of skin bacteria, such as *S. epidermidis*^10,32^, might negatively impact tick health. To test this, we compared feeding metrics of *dae2*^*Is*^ and control ticks at day 3 by first quantifying tick body expansion across two axes. The ratio of these measurements, known as the scutal index (SI), reflects how long ticks have been feeding on a bloodmeal host (Fig. 5c)^33^. Control ticks had significantly higher SIs than *dae2*^*Is*^ knockdown ticks (Fig. 5d). We also weighed individual ticks and found that *dae2*^*Is*^ ticks were lighter than controls (Fig. 5e), suggesting increased host bacteria interfered with feeding. Impaired feeding may be due to slowed attachment, reduced blood intake, or some combination of these, ultimately resulting in decreased feeding or more prolonged, risky interactions with a bloodmeal host. In total, our combined *in vitro* and *in vivo* analyses reveal that Dae2^*Is*^ limits levels of Gram-positive host skin commensals, such as *S. epidermidis*, enhancing the ability of *I. scapularis* to resist against damaging infections by pathogens confronted during blood-feeding. Taken together, these data suggest that Dae2^*Is*^ mediates resistance to host skin commensals, most likely by inhibiting bacteria both at the tick-host feeding site and within the tick gut.

Expansion of antimicrobial strategies was likely a key facilitator for the unique lifestyle of hematophagous arthropods. The rise of blood-feeding behavior over the course of arthropod evolution was concomitant with the emergence of numerous antibacterial genes, such as *dae2*^34^. The intriguing timing of *dae2* acquisition by ticks and mites soon after the emergence of hematophagy within this arthropod clade might indicate that Dae2 activity was an important enabler of blood-feeding^2^. Hard ticks imbibe blood continuously from one host for several days at a time, leading to exceptionally intimate and sustained interactions with a wide range of vertebrates and associated microbes^35^. The possibility that Dae2^*Is*^ could act in both the gut and saliva of ticks would suggest that immune function both in- and outside of the organism could be of great biological importance to ectoparasites, such as ticks. Antimicrobial activity in tick saliva could also provide an additional benefit of blocking contamination of the bite site itself, preventing a cascade of antagonistic host responses within the dermis.

More broadly, our results underscore how relationships between animals and bacteria are context-dependent^36^. Pathogenicity of a given microbe refers to a status, rather than a strict identity. Eukaryotes have evolved to respond appropriately; thus, the differential specificity of animal immune systems reflects how host–microbe interactions vary between organisms. Ticks harbor symbiotic microbes, such as *B. burgdorferi*, which are tolerated by their tick vector but are pathogenic to humans upon transmission^28^. In this study, we discovered that the mirror opposite is also true: ticks are vulnerable to infection by bacterial commensals found on the skin of vertebrates they must feed on. Finally, our observation that *dae2*^*Is*^ has evolved to target bacterial partners of vertebrate hosts highlights both the inseparability of animal-microbiome relationships and also the transmutable nature of microbial interactions across different biological contexts. Animals not only confront each other but also with associated microbes, which are an integral part of an organismal “package.” Particularly for blood-feeding parasites, such as ticks, successful navigation of these complex, interkingdom clashes is critical for survival.

## Acknowledgements

We are grateful to George Meigs and James Holton at the Advance Light Source (Lawrence Berkeley National Laboratory), the Stroud lab (UCSF) and Michael Thompson (Fraser lab, UCSF) for help with X-ray crystallography. We thank Carol Gross, Tiffany Scharschmidt and Joe Bondy-Denomy (all at UCSF) as well as Patricia Rosa (Rocky Mountain Laboratories, NIAID, NIH) for providing bacterial strains. We also appreciate Dyche Mullins and Samuel Lord (UCSF) for their assistance with microscopy. The Chou lab thanks Soraya Pedemonte and Ethel Enoex-Godonoo for lab and administrative help, respectively, without which the lab would not run as smoothly. We are also grateful to Harmit Malik (UCSF), Carol Gross (UCSF), Wallace Marshall (UCSF), Tiffany Scharschmidt (UCSF), Jeremy Reiter (UCSF), Thea Mauro (UCSF), Sophie Dumont (UCSF), and the entire Chou lab for helpful discussions and review of the manuscript. This project was funded in part by NIH grants (R01AI132851 to SC, R01AI134696 to JP), Research Councils UK (EP/T002778/1) to WV, and UCSF Program for Breakthrough Biomedical Research, funded in part by the Sandler Foundation (to SC). Additional support for SC came from the Chan Zuckerberg Biohub the Johnson & Johnson WiSTEM2D Award. FY was supported by the National Science Foundation (1650113) and a grant to UCSF from the Howard Hughes Medical Institute through the James H. Gilliam Fellowships for Advanced Study program.

## Author Contributions

B.M.H., A.D.R., J.H.F.P, and S.C. designed the study. B.M.H., A.D.R., F.Y., S.F., J.K., Z.Z., K.W.L., L.M., J.B., and V.B. performed experiments. All authors analyzed data and provided intellectual input into aspects of this study. B.M.H, A.D.R., J.H.F.P., and S.C. wrote the manuscript; all authors contributed to its editing.

