## Extended Data and Material and Methods for "Immune factor of bacterial origin protects ticks against host skin microbes"

### Extended Data and Materials and Methods

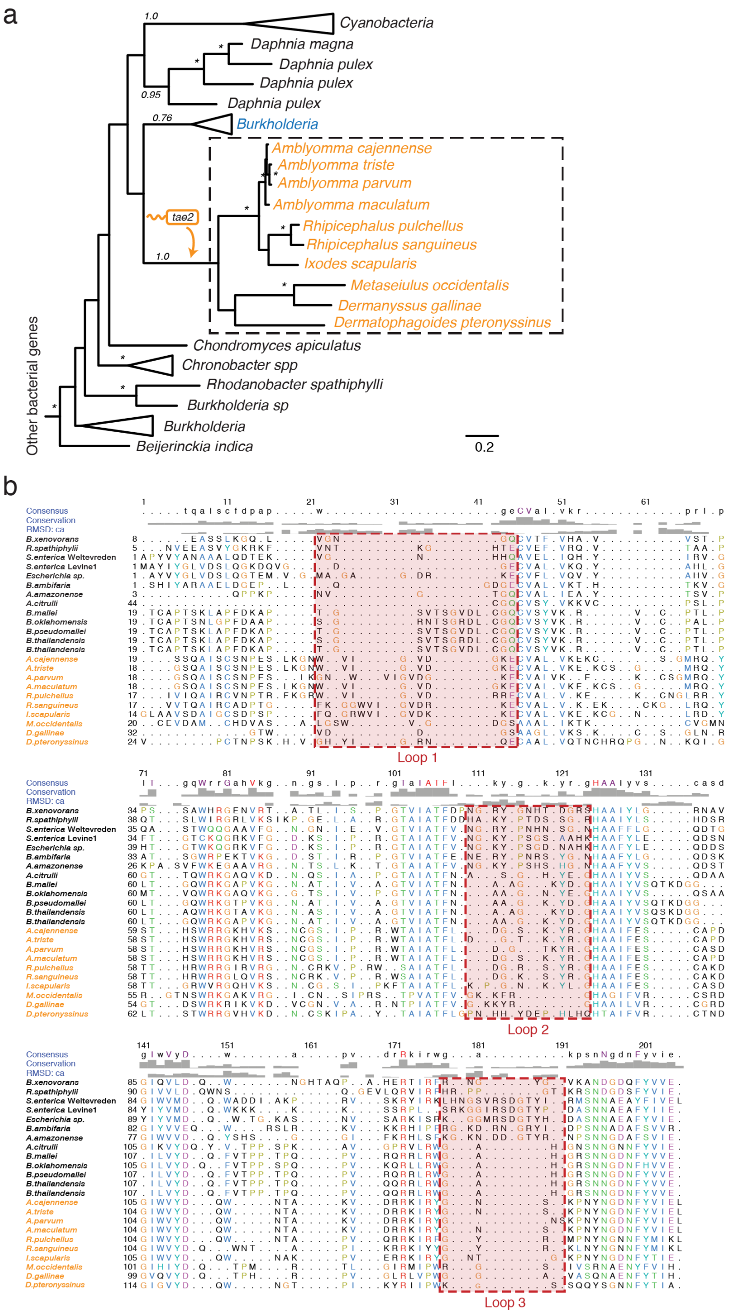

**Figure 1 | Evolutionary analysis of Tae2$^{St}$ and Dae2$^{Is}$ enzymes. a,** Phylogenetic tree demonstrates horizontal transfer of genes encoding Tae2 into a select group of eukaryotic organisms, including Deer ticks, Lone star ticks, Brown dog ticks, as well as mites. **b,** Protein sequence alignment of bioinformatically identified Tae2 and Dae2 enzymes. The bacterial Tae2 enzymes are shown in black letters, while their eukaryotic Dae2 counterparts are in orange letters. The boxed regions (red boxes) correspond to highly variable structural loop elements. Loops correspond to those denoted in main text Fig. 1b.

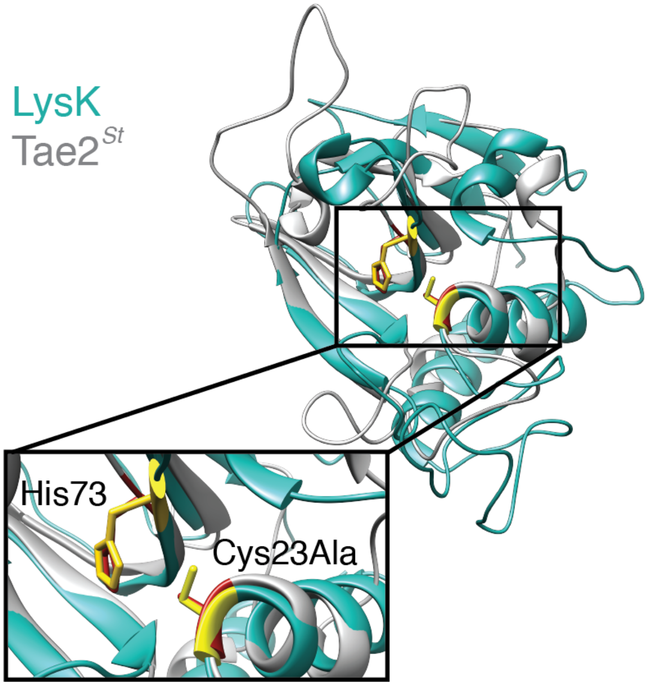

**Figure 2 | Structural alignment of Tae2$^{St}$ with its closest structural homolog, LysK from staphylococcal phage K (PDB ID: 4CSH).** High conservation was observed regarding the catalytic residues (Ala23 and His73), which is contrast to multiple differences in helices and loops throughout the proteins. Although Tae2$^{St}$ does adopt a known amidase fold, this high degree of variability (RMSD 3.0 and Percent identity 14%) suggests a potential unique functional role of this enzyme relative to known structural homologs.

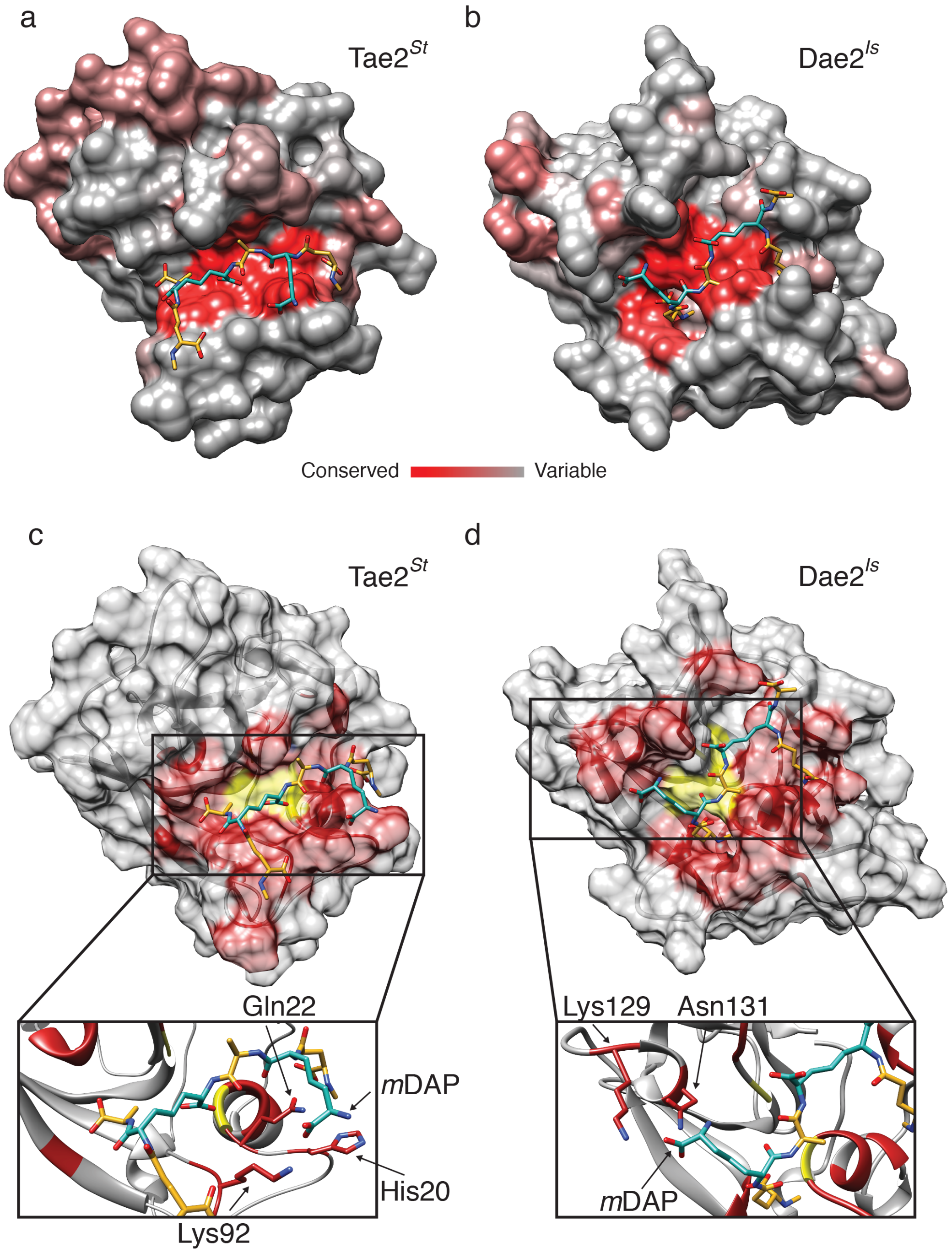

**Figure 3 | Active site conservation and substrate docking analysis. a, b,** Consurf server analysis indicates that the majority of conservation in the case of both enzymes, Tae2$^{St}$ (a) and Dae2$^{Is}$ (b) is within the active site. Highly conserved residues are shown in red, and least conserved in grey. The PG substrate for each is also included to further highlight the active site. **c,** Tae2$^{St}$ and **d,** Dae2$^{Is}$ residues interacting with the PG substrate are highlighted in red. These were determined based on our docking analysis. The close-up insets demonstrate the enzyme residues coordinating the same *m*DAP residue in the PG substrate.

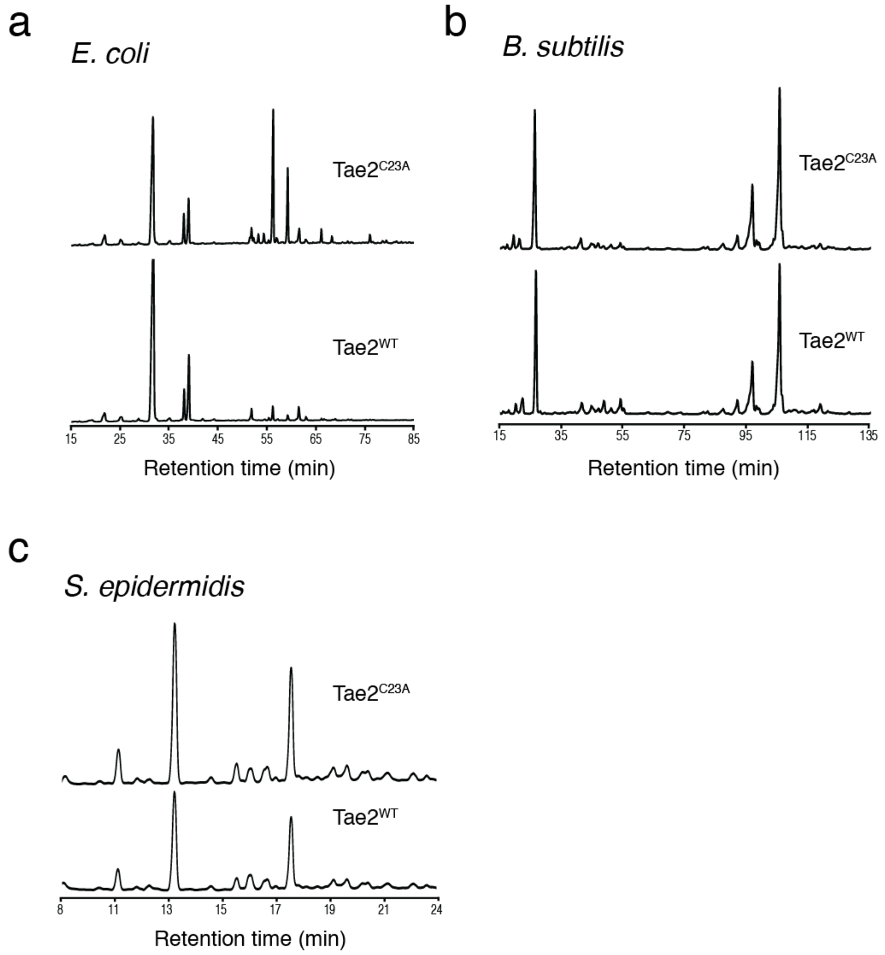

**Figure 4 | Analysis of the enzymatic ability of Tae2$^{St}$ to digest PG substrates. a,** HPLC chromatograms demonstrate the disappearance of *E. coli* PG fragments in the assays with Tae2$^{St}$ WT, but not with the catalytically inactive Tae2$^{St}$ C23A. Tae2$^{St}$ WT does not degrade PG substrates from either of two Gram-positive PG substrates **b,** *B. subtilis* and **c,** *S. epidermidis*.

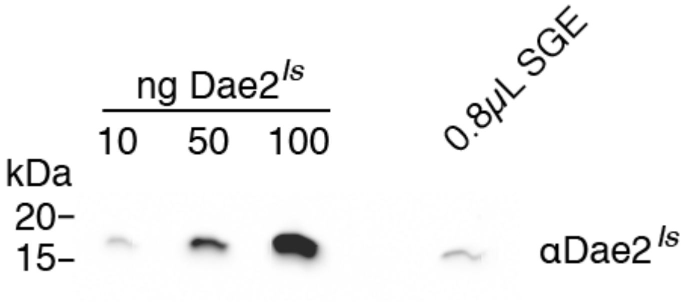

**Figure 5 | Quantification of Dae2$^{Is}$ in salivary glands.** Densitometry was used to estimate the concentration of Dae2$^{Is}$ in *I. scapularis* salivary gland extract (SGE). Known amounts of recombinant Dae2$^{Is}$ (10, 50, and 100 ng) were stained with αDae2$^{Is}$ and a secondary anti-rabbit to create a standard curve. Intensities were determined with ImageJ software and compared to that of SGE.

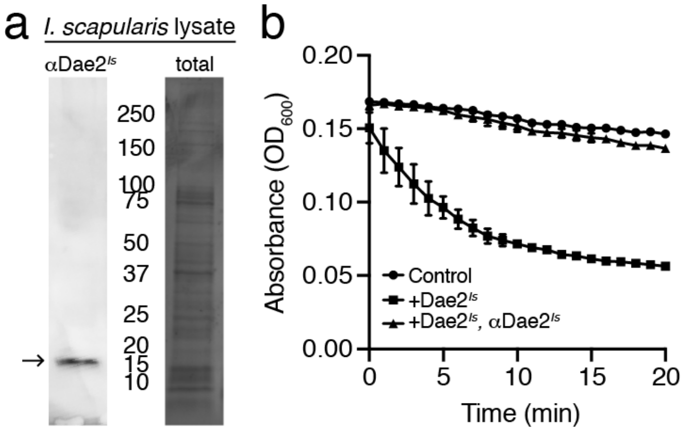

**Figure 6 | αDae2$^{Is}$ specifically blocks Dae2$^{Is}$ enzyme activity. a,** Western blot analysis of total *I. scapularis* female tick homogenate using αDae2$^{Is}$. Single band resulting from blot appears at predicted size of Dae2$^{Is}$ and is labeled (left, arrow). An equivalent load-control coomassie blue-stained gel is also shown (right). **b,** Dae2$^{Is}$ lysis assays against *Bacillus subtilis* cells. Recombinant Dae2$^{Is}$ was added to log-phase *B. subtilis* cells at 10μM +/- αDae2$^{Is}$ at 1:1 molar ratio, and cell viability was measured over time. Buffered cells without enzyme were included as a control.

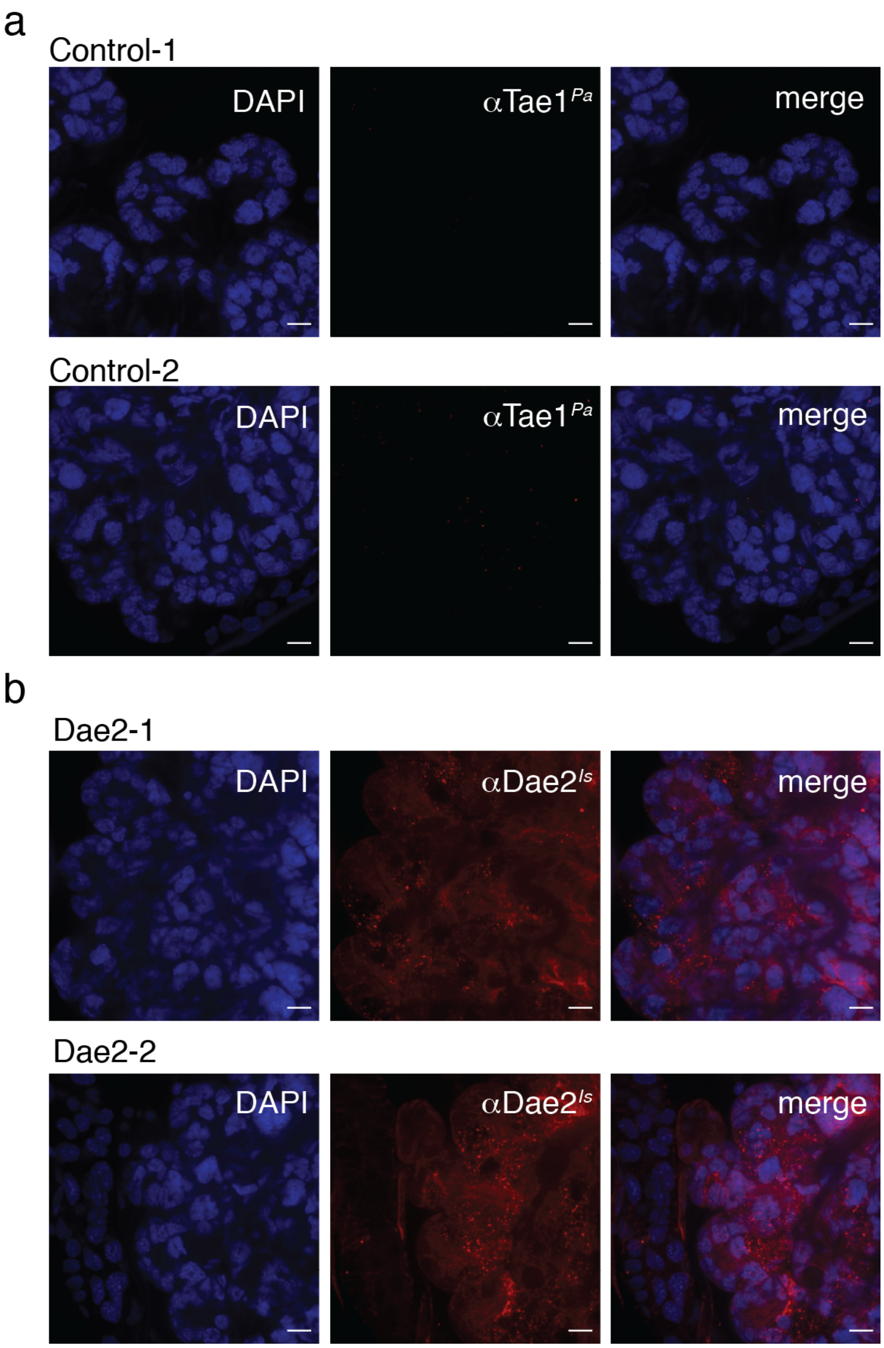

**Figure 7 | Dae2$^{Is}$ localization in tick salivary glands** Additional confocal images of *I. scapularis* salivary glands from adult female ticks. Whole mounts were stained with DAPI (left panels, blue), control rabbit antibody against Tae1$^{Pa}$ (**a**, middle, red) or rabbit αDae2$^{Is}$ (**b**, middle, red). Merged images shown on the right. Scale bars= 15µm.

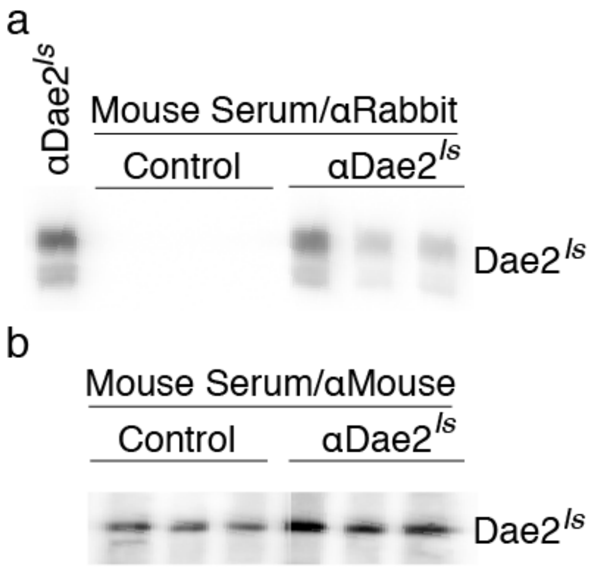

**Figure 8 | αDae2$^{Is}$ stability in immunized and tick-infested mice. a,** Western blot analysis against recombinant Dae2$^{Is}$ using α-rabbit secondary antibody against serum collected from control (left) and αDae2$^{Is}$-immunized (right) mice at 18 days following antibody injection in experiment diagramed in main text Fig. 5. αDae2$^{Is}$ was generated in rabbit; thus α-rabbit signal should be specific for the injected αDae2$^{Is}$ antibodies. αDae2$^{Is}$ was used as a control (far left). **b,** α-mouse secondary was used to detect primary mouse antibodies made against Dae2$^{Is}$. Both groups develop similar mouse antibodies against Dae2$^{Is}$ after *I. scapularis* tick feeding.

**Table 1 | Data collection and refinement statistics for Tae2$^{St}$.**

| | Crystal 1 |
|---|---|
| **Data collection** | |
| Space group | P 1 21 1 |
| Cell dimensions | |
| *a*, *b*, *c* (Å) | 33.318 46.032 38.923 |
| Resolution (Å) | 37.65 - 2.05 (2.123 - 2.05) |
| $R_{sym}$ or $R_{merge}$ | 0.0623 (0.1245) |
| *I* / *I* | 19.94 (7.39) |
| Completeness (%) | 98.21 (86.31) |
| Redundancy | 1.99 (1.97) |
| **Refinement** | |
| Resolution (Å) | 37.65 - 2.05 |
| No. reflections | 7120 (624) |
| $R_{work}$ / $R_{free}$ | 0.1752 / 0.2223 |
| No. atoms | 1196 |
| Protein | 1109 |
| Water | 87 |
| *B*-factors | |
| Protein | 23.55 |
| Water | 27.69 |
| R.m.s. deviations | |
| Bond lengths (Å) | 0.007 |
| Bond angles (°) | 0.81 |

*Values in parentheses are for highest-resolution shell.

**Table 2 | Primers used in this study.**

| | Name | Sequence 5'->3' | Reference (PMID) |
|---|---|---|---|
| **PCR** | | | |
| | Tae2$^{St}$-C23A-QC-For | CCACCATCAGGCTGTTGAACTTATTCAACACTATA | this paper |
| | Tae2$^{St}$-C23A-QC-Rev | TAAGTTCAACAGCCTGATGGTGGTTTCCTACCTTT | this paper |
| **qPCR** | | | |
| | Is_actin_For | ACCTGACCGACTACCTGATG | 25470067 |
| | Is_actin_Rev | CAGAGCTTCTCCTTGATGTCG | 25470067 |
| | Is_dae2_For | GAAGCTTTTTCTCATCAGCGCAG | 25470067 |
| | Is_dae2_Rev | CAATTCGATCGTGTAGAAGTTGTC | 25470067 |
| | Staph_genus_For (TStaG422) | GGCCGTGTTGAACGTGGTCAAATCA | 11427566 |
| | Staph_genus_Rev (TStag765) | TIACCATTTCAGTACCTTCTGGTAA | 11427566 |
| **siRNA** | | | |
| | dae2Is_start310_antisense | AAGTACTTGGGTCACGCCGCCCCTGTCTC | this paper |
| | dae2Is_start301_sense | AAGGCGGCGTGACCCAAGTACCCTGTCTC | this paper |
| | dae2Is_start281_sense | AAGCTTGAGGAAAGTGGCGATCCTGTCTC | this paper |
| | dae2Is_start281_antisense | CCATCGCCACTTTCCTCAAGCCCTGTCTC | this paper |
| | dae2Is_start375_antisense | AAATACAGGGAAGCCCAACTACCTGTCTC | this paper |
| | dae2Is_start375_sense | AATAGTTGGGCTTCCCTGTATCCTGTCTC | this paper |
| | dae2Is_scrmble281_sense | ATGAAGAGAAGGTTGTACCGGCCTGTCTC | this paper |
| | dae2Is_scramble281_antisense | GCCCTCCTCCCGCTCATATAACCTGTCTC | this paper |
| | dae2Is_scrmble375_antisense | ACGAAGCATCCGAGAATAACACCTGTCTC | this paper |
| | dae2Is_scrmble375_sense | AATTTACTCTGTGCGTGACGTCCTGTCTC | this paper |
| | dae2Is_scramble301_sense | AACGCCGACTATGACGCAGGCGCTGTCTC | this paper |
| | dae2Is_scramble301_antisense | AAGCCTCATCCGGCGTATCGGCCTGTCTC | this paper |

#### Materials and Methods

##### Cloning, expression, and purification of Tae2 and Dae2 enzymes

Full-length Tae2 from Salmonella typhi (Tae2$^{St}$) in the pET29b+ expression vector has been described previously[1]. To create the catalytically dead mutant (C23A), we used a quick-change protocol employing Tae2$^{St}$-C23A-QC-Forward and reverse primers (Extended Data Table 2) followed by DpnI digestion before transforming into chemically competent cells (Macrolab). Luria Broth (LB) was used for routine cloning with DH5alpha or XL1Blues and proteins were expressed and purified from BL21 or Rosetta2 (DE3) *E. coli* strains grown in Terrific Broth (TB).

Protein expression was induced at an optical density at 600 nm ($OD_{600}$) of 0.6 by the addition of 1.0 mM IPTG. Tae2 was induced for 3 hrs at 37 °C and lysed by sonication. Cell lysate was incubated with metal-chelating affinity resin, followed by resin washes, and final protein elution with the same buffer (except using 400 mM imidazole). The protein was further purified via size exclusion chromatography (GE Healthcare). For Dae2$^{Is}$, *E. coli* was grown to an $OD_{600}$ of 0.6, followed by cell resuspension in 20 mM HEPES pH 7.5, 0.1 M NaCl, 25 mM imidazole. Cells were then induced with 1.0 mM IPTG overnight at 16 °C. The protein was further purified via a metal-chelating affinity column and size-exclusion chromatography (GE healthcare) as described above.

To prepare selenomethionine (SeMet; Sigma-Aldrich) protein derivatives for determining crystal structure, proteins were purified as above but with the follog differences. *E. coli* cells were grown in M9 minimal media to an $OD_{600}$ of 0.6 before 60 mg SeMet, 100 mg each of threonine, lysine hydrochloride, and phenylalanine, and 50 mg each of leucine, isoleucine, and valine, were added as solids[2]. The culture was incubated for 15 min before inducing with 1.0 mM IPTG.

##### Protein structure determination and analysis

Tae2 C23A crystals were generated by hanging drop vapor diffusion at 25 °C from a 1:1 mixture of 4.32 mg/ml protein in 20 mM HEPES pH 7.5, 0.1 M NaCl, 0.02 % Na azide with 0.1 M MES pH 6.6, 1 M sodium citrate, and 4 % formamide for 24 hrs. Crystals were directly used for diffraction data collection at the Lawrence Berkeley National Laboratory Advanced Light Source Beamline 8.3.1. Phases were obtained experimentally with data from a SeMet-substituted crystal with the PHENIX software suite[3]. Coot[4] and maximum likelihood refinement with PHENIX[3] were used for iterative building and refinement. Coordinates and structural factors were deposited in the Protein Data Bank (PDB ID: 6WIN). The closest structural homolog (4CSH) identified through the DALI server[5] was visualized with Chimera[6]. Structural models of Tae2 family homologs, as well as Dae2 family homologs, were built with the PHYRE One-to-One threading algorithm using the default homology modeling parameters, including a local alignment and weighted secondary structure scoring (weight of 0.1) [7]. Structures were aligned using Chimera[6].

##### Ligand model generation

Initial protein preparation commenced from the crystal structure of Tae2$^{St}$ C23A. The mutation of the catalytic residue Cys23 to Ala for crystalline stability was reverted back to wild-type Cys. This 'WT' crystal structure was used as the input for building a homology model of

Dae2$^{Is}$. For initial structural studies using Molecular Operating Environment (MOE) 2018 (Chemical Computing Group), protonation states were assigned with MOE2018. Later structural studies were performed using Maestro (Schrodinger) and these systems were prepared by the Protein Preparation Wizard. Maestro's Binding Surface Area Analysis tool was used to compare the available binding surface between Tae2$^{St}$ and Dae2$^{Is}$ in the context of the docked conformations (see below).

**Computational analysis of docking poses**

Given the conformational complexity of a cross-linked peptidoglycan (PG), and the resultant sampling problem that occurs when trying to dock this highly flexible compound to a shallow binding surface, we used crystal structures of a single PG, MLD (PDB IDs: 2CB3, 2F2L, 4QRB, 4QR7, 4QTF, 6I9O) to form guesses at the cross-linked PG conformational space. Conformational space for these PGs was sampled using the Conformational Search algorithm within MOE, and all moderate-to-low-energy (50 kcal/mol cut-off) conformations were retained for docking. Initial rigid docking of these linked fragments to Tae2$^{St}$ and Dae2$^{Is}$ was performed with MOE2018, using the AMBER10:EHT forcefield and default options for scoring and returning poses. Based on these results, we elected to truncate the PG to the cross-linked tripeptide dimer as a means of simplifying the search space. We present results wherein the cross-linked peptidoglycan has been truncated to a simplified representation of the true substrate; two tripeptide stems consisting of D-glutamic acid (D-Glu)—meso-diaminopimelic acid (*m*DAP)—D-Alanine (D-Ala), with an amide bond (cross-link) between D-Ala in one stem and *m*DAP in the other stem.

**Peptidoglycan purification, analysis, and enzyme assays**

Peptidoglycan (PG) was purified as previously described[8,9]. Bacterial strains were grown to an $OD_{600}$ of 0.6, harvested by centrifugation and boiled in SDS (4 % final concentration) for 4 hours with stirring. After washing in purified water to remove SDS, the peptidoglycan was treated with Pronase E for 2 h at 60°C (0.1 mg/ml final concentration in 10 mM Tris-HCl pH7.2 and 0.06 % NaCl; pre-activated for 2 h at 60 °C). Pronase E was heat inactivated at 100 °C for 10 min and washed with sterile filtered water (5 x 20 min at 21k × *g*). PG from Gram-positive bacteria was purified as described above and treated with 48% HF at 37°C for 48 h to remove teichoic acids, followed by washes with sterile filtered water before Tae2$^{St}$ and Dae2$^{Is}$ enzyme degradation (0.1-1 µM, 4 h at 37°C in 10 mM Tris-HCl pH 7.2 and 0.06% NaCl). Enzymes were heat inactivated at 100°C for 10 min. Mutanolysin (Sigma M9901, final concentration 20 µg/ml) or cellosyl (kindly provided by Hoechst, Frankfurt Germany) was directly added to the purified peptidoglycan and incubated overnight at 37 °C. The peptidoglycan fragments were reduced, acidified, analyzed via HPLC as described previously[8,9]. Alternatively, a modified HPLC method (0.5 ml/min flow rate, 55°C with Hypersil ODS C18 HPLC column, Thermo Scientific, catalog number: 30103-254630) was used: start at 100 % buffer A, ramp to 100 % buffer B over 20 min, maintain 100 % buffer B for 2 min, ramp to 100 % buffer A over 2 min, maintain 100% buffer A over 30 min.

**Tick feeding**

*I. scapularis* nymphs and adults were purchased from the Tick Lab at Oklahoma State University (OSU) or provided by BEI Resources, a division of the Center for Disease Control.

Ticks were maintained in glass jars with a relative humidity of 95% (saturated solution of potassium nitrate) in a sealed incubator at 22°C with a light cycle of 16h/8h (light/dark). Animal experiments were conducted in accordance with the approval of the Institutional Animal Care and Use Committee (IACUC) at UCSF. Ticks were fed on young female C3H/HeJ mice acquired from Jackson Laboratories. Mice were anesthetized with ketamine/xylazine before placing ≤30 nymphs or ≤6 adult female ticks. Ticks were either pulled off of isoflurance anesthetized mice at various times during feeding (1-7 days) or allowed to feed to repletion and collected from mouse cages.

##### Tick saliva collection

Partially fed (5-7 days) female ticks were taped to glass slides. Saliva production was stimulated by pipetting 2-5μL of 5% pilocarpine hydrochloride (Tokyo Chemical Industries) in methanol onto the scutum. Saliva was collected in capillary tubes fitted onto the hypostome (mouthparts) of the ticks during a period of 24 hours in a humid chamber. Saliva was stored at -20°C before SDS-PAGE analysis.

##### Mouse serum collection

One to two weeks following tick feeding, blood was collected from mice. Whole blood was centrifuged at 7k x g for 8 minutes at 4°C. Serum (top layer) was transferred to new tubes and stored at -20° until use.

##### SDS-PAGE and Western Blotting

Proteins were separated by electrophoresis and transferred to a nitrocellulose membrane with a BioRad Trans-Blot Turbo system. Membranes were blocked in 5% nonfat milk in TBST (Tris-buffered Saline 0.1% Tween20). Rabbit αDae2$^{Is}$[10] was used at 1:2000 in TBST followed by Goat αRabbit-HRP (Advansta R-05072-500) at 1:5000 in TBST. Chemiluminescence was detected using Biorad Clarity Western ECL substrates and imaged on an Azure c400 instrument. For tick saliva western blot, recombinant Dae2$^{Is}$ was used as the positive control. For mouse serum reactivity to Dae2$^{Is}$, Dae2$^{Is}$ was loaded into a single well 15% acrylamide gel, transferred to a membrane and blocked as described above. Reactivity with Rabbit αDae2$^{Is}$ antibody was used as a positive control. Mouse serum was diluted 1:200 in TBST followed by Goat αMouse-HRP (Advansta R-05071-500) at 1:5000.

##### Microscopy

Salivary glands dissected from adult female *I. scapularis* were fixed in formalin for 3 days at room temperature. After washing 3 times in PBST (PBS with 0.3% Triton X-100) for 15 minutes at 37°C, salivary glands were incubated in blocking buffer (PBST with 10% fetal bovine serum, Sigma) for 30 minutes at 37°C[10] . Primary antibodies, αDae2$^{Is}$ and control αTae1[10] were diluted at 1:500 in blocking buffer and incubated overnight at 4°C. After washing 3 times in PBST, donkey αRabbit Alexa Fluor 647 (Invitrogen) diluted 1:1000 in the blocking buffer was added for 1 hour at room temperature in the dark. Samples were washed 3 times in PBST and stained with DAPI (Invitrogen, 50ng/ml in PBST) for 30 minutes at room temperature in the dark. After washing 3 times in PBST salivary glands were mounted on microscope slides with 10 µl ProLong Diamond Antifade Mounting media (Invitrogen). Fluorescence imaging was performed on a Nikon spinning disk confocal microscope using a 60x/1.49 objective.

**Tick RNA isolation and qRT-PCR**

RNA and DNA from ticks was isolated using Trizol reagent (Fisher Scientific) according to the manufacturer's protocol. *I. scapularis dae2* and *actin* transcripts were quantified with RT-PCR as described previously[10]. Primers specific to the staphylococcus genus were used to quantify bacterial load in tick DNA samples using the Applied Biosystems PowerUp SYBR Green Master Mix. All primers are listed in Supplementary Table 2.

**Bacterial Growth**

*Borrelia burgdorferi* strain S9[11] was provided by Dr. Patricia Rosa (NIAID, NIH, RML) and cultured in BSK II media at 35°C, 2.5% $CO_2$. *Staphylococcus epidermidis* (BCM060 and SK135), *S. hominis* (SK119), *Corynebacterium propinquum* (DSM44285), and *C. jeikeium* (DSM7171) isolates were provided by Dr. Tiffany Scharschmidt (UCSF). *Staphylococcus* species were grown in Tryptic Soy Broth at 37° C with shaking. *Corynebacterium* were cultured in Blood Heart Infusion (BHI) supplemented with 10% Tween 80. *Bacillus subtilis* (168) was provided by Dr. Carol Gross and cultured in LB at 37° C with shaking.

**Lysis Assays**

Log-phase bacteria were incubated with $Dae2^{Is}$ or $Tae2^{St}$ (wt or catalytic mutant) at 0.5-5µM in low salt buffer (20mM Hepes pH 7.0, 100mM NaCl) for 4 hours. The reaction was plated on appropriate solid media (see above) and incubated to determine CFU. Results are presented as percent survival (CFU surviving in wt normalized to its respective catalytic mutant).

**Tick Injection/Viability Assays**

Log-phase bacteria were diluted in PBS to a concentration of $10^7$ CFU/mL and injected into *I. scapularis* nymphs using an Eppendorf FemtoJet 4x microinjector. At varying times post injection, ticks were crushed in PBS and plated on appropriate solid media to determine CFU. Injections of PBS alone was used as a monitor of tick fitness and survival of the injection trauma. To assess tick viability, ticks were monitored every hour after injection for movement and reaction to $CO_2$ (human breath). Ticks that had curled their front legs, did not respond to $CO_2$, and were not mobile even after picking up with forceps (no reflex straightening of legs) were considered dead.

**Pre-immunization of mice against Dae2**

24 hours before tick feeding, 0.5mg affinity purified Rabbit αDae2 custom generated against recombinant $Dae2^{Is}$ protein (Genscript) was diluted in sterile PBS and injected into the intraperitoneal cavity of female C3H/HeJ mice to neutralize against pre-made $Dae2^{Is}$ protein in tick saliva. Sterile PBS was injected into control mice.

**RNAi knockdown of *dae2* in feeding nymphs**

*dae2$^{Is}$* targeting and scrambled primers listed in Extended Data Table 2 were used with the Ambion Silencer siRNA Construction Kit (AM1620) to generate three different siRNA constructs targeting *dae2*$^{Is}$, or equivalent scramble controls. Constructs were pooled such that each siRNA had a final concentration of 600 ng/µl. Five microliters of the siRNA pool was loaded into a capillary tube and injected into the midgut of *I. scapularis* nymphs. Microinjected nymphs were allowed to recover overnight in a humid environment. Nymphs were fed on mice pre-immunized with αDae2 (see above) and forcibly removed from mice after 3 days. Ticks were weighed and imaged to determine scutal indices. RNA was isolated from pools of 3 ticks (ordered by increasing scutal index) and *dae2*$^{Is}$ KD efficiency was determined by qRT-PCR as described above.
